# Elucidating redox-driven inhibition of methanogenesis by an artificial quinone in *Methanosarcina barkeri*: Integrated proteomic and physiological evidence

**DOI:** 10.64898/2026.01.27.701717

**Authors:** Paola A. Palacios, Hugo Kleikamp, Jeppe L. Nielsen, Mathias Eskildsen, Anders Bentien, Michael V. W. Kofoed

## Abstract

Methanogenesis is a crucial component of Earth’s carbon cycle and a source of methane for biofuel production. The presence of higher energy electron acceptors, such as iron(III) oxides and quinones, is believed to significantly impact methanogenesis. This study investigated the physiological and proteomic responses of the type I *Methanosarcina, M. barkeri*, to the artificial quinone 9,10-anthraquinone-2,7-disulfonate disodium (2,7-AQDS), using H_2_/CO_2_ as substrates. Our findings revealed that during 2,7-AQDS reduction, cellular growth ceased. The lack of energy conservation was associated with direct inhibition of both methanogenesis and CO_2_ utilization, corroborated by a significant downregulation of the enzymes involved in this metabolic pathway. Furthermore, the significant upregulation of specific subunits of the reversible Ech hydrogenase suggests that this enzyme redirects electrons from H_2_ towards the most energetically favorable reaction (2,7-AQDS reduction), rather than the reduction of ferredoxin, which is a highly energy-demanding process, essential for initiating the CO_2_ reduction pathway. Additionally, it is conceivable that Ech homologues in other hydrogenotrophic methanogens also participate in the reduction of higher energy-yielding electron acceptors. These findings provide novel insights into how quinones, particularly in their oxidized state, directly impact methanogenesis, thereby influencing both artificial and natural methanogenic environments.

## 1. INTRODUCTION

Methanogenesis, a pivotal process in the Earth’s carbon cycle, is primarily carried out by methanogenic Archaea. These microorganisms efficiently conserve energy for ATP synthesis through the production of methane (Balch et al. 1979). Thereby, methanogens contribute substantially to the release of methane to the atmosphere (Buan 2018), directly impacting climate change and global warming (Buan 2018). Also, they play a crucial role in the formation of natural gas reserves (Deppenmeier 2002), which are vital for domestic or industrial energy needs such as heating, electricity generation, and transportation.

Methanogens inhabit a diverse range of anoxic environments, including marine and freshwater sediments, flooded soils, gastrointestinal tracts, anaerobic digestors, landfills, geothermal systems, and extreme environments (Liu and Whitman 2008; Buan 2018). While methanogenesis is a strictly anaerobic process (Lyu et al. 2018), certain systems like anaerobic digestors, biomethanation reactors, and gastrointestinal tracts are prone to oxygen exposure. Research indicates that some methanogens have developed various strategies to endure oxygen stress (Lyu and Lu 2017; Kleikamp et al. 2023), an area of ongoing investigation.

In the biogeochemical redox chain, oxygen serves as the primary electron acceptor due to its high redox potential. In anoxic environments, the redox chain continues with nitrate, manganese(IV)oxide, iron(III)oxides, sulfate and sulfur and, finally, with CO_2_ (Schink 2006). This means that electrons derived from organic matter oxidation run preferentially to the electron acceptor with the higher redox potential, yielding the highest amount of energy. Each redox process, influenced by the type of electron acceptor available, is catalyzed by specific microbial groups that contribute to the different biogeochemical cycles. Hence, under conditions with low redox potential where alternative electron acceptors to CO_2_ are scarce, methanogenesis and CO_2_ reduction become favorable (Lyu et al. 2018).

However, in both natural and engineered anoxic environments, methanogens encounter higher energy-yielding electron acceptors, such as iron(III) oxides and quinones (Zhou et al. 2014). Quinones are naturally present in soils, comprising ∼60% of organic matter, and in sediments, in the form of humic substances (HS) (Cervantes et al. 2000; Trevisan et al. 2010; Xu et al. 2013), a recalcitrant but redox active mixture of phenolic carboxylic acids derived from the biochemical degradation of plant and animal residues (Muscolo et al. 2013; Zavarzina et al. 2021). Due to their redox nature, electron transfer to and from HS is fully reversible over successive redox cycles, and previous research has suggested that this may largely suppress methane production particularly in temporarily anoxic systems such as wetlands, which largely impact the global methane flux (Klüpfel et al. 2014). Beyond their ecological role in reduction of methane production (Xu et al. 2013; Klüpfel et al. 2014; Kulikova and Perminova 2021), quinones have also been explored for biotechnological purposes (Cervantes et al. 2003; Liang et al. 2020; Xu et al. 2022; Palacios et al. 2023; Tucci et al. 2023). For instance, the reduction of anthraquinone-2,6-disulfonate (2,6-AQDS), a humic substance analog, has been evidenced in different methanogens including *Methanosarcina acetivorans* WWM1, *Methanosarcina barkeri* MS, *Methanosphaera cuniculi* 1R7, *Methanobacterium palustre* F, and *Methanococcus voltaei* A3 (Bond and Lovley 2002; Holmes et al. 2019; Eliani-Russak et al. 2023). Except for the non-hydrogenotrophic *M. acetivorans*, which can conserve energy via an exclusive membrane-bound multiheme *c*-type cytochrome (MmcA) during methanol utilization and 2,6-AQDS reduction (Holmes et al. 2019), AQDS mediated iron reduction by the other hydrogen-utilizing methanogens resulted in growth and methanogenesis inhibition (Bond and Lovley 2002).

Consequently, beyond *M. acetivorans*, our understanding of how other *Methanosarcinales* or methanogens, with a distinct set of enzymes for energy conservation and without multiheme *c*-type cytochromes, are impacted by the presence of quinones and catalyze their reduction remains an open research question. Given the metabolic versatility of *M. barkeri*, capable of performing hydrogenotrophic, acetoclastic, and methylotrophic methanogenesis, as well as their ubiquity in both natural and artificial methanogenic habitats (Bryant and Boone 1987; He et al. 2019; Zhou et al. 2021), *M. barkeri* was selected as a model organism for this study. This research investigates the effects of 9,10-anthraquinone-2,7-disulfonate disodium (2,7-AQDS) on methanogenesis, and cell growth, while also exploring the proteomic responses of *M. barkeri* during the reduction of this higher-energy yielding electron acceptor. Ultimately, by obtaining a deeper understanding on the mechanisms behind quinone reduction and its impact in methanogenesis, this study seeks to shed light on the ecological and biotechnological implications of quinone exposure, particularly in their oxidized form.

## 2. MATERIALS AND METHODS

### 2.1. Culture conditions

*Methanosarcina barkeri* strain 227 (DSM 1538) was obtained from the German collection of microorganisms and cell cultures DSMZ (https://www.dsmz.de). Subcultures of *M. barkeri* were sustained in a modified DSMZ medium 120 containing K_2_HPO_4_ (0.35 g L^-1^), KH_2_PO_4_ (0.23 g L^-1^), NH_4_Cl (0.50 g L^-1^), MgSO_4_ × 7H_2_O (0.50 g L^-1^), CaCl_2_ × 2H_2_O (0.25 g L^-1^), NaCl (2.25 g L^-1^), FeSO_4_ × 7H_2_O solution (0.1% w/v in 0.1 H_2_SO_4_) (2 mL L^-1^), trace element solution SL-10 (320 DSMZ medium) (1 mL L^-1^), Na-resazurin solution (0.1% w/v) (0.5 mL L^-1^), NaHCO_3_ (0.85 g L^-1^), L-cysteine-HCl x H_2_O (0.3 g L^-1^), and vitamin solution (141 DSMZ medium) (10 mL L^-1^). Cultures were maintained in butyl-rubber-stoppered glass bottles with an anoxic headspace of H_2_/CO_2_ (80:20, v/v) at 1.7 bar and 37 °C.

### 2.2. Experimental design

Three conditions were monitored to assess the reduction of AQDS by *M. barkeri*: (i) AQDS supplementation under abiotic conditions, (ii) AQDS supplementation using *M. barkeri* as a biocatalyst, and (iii) a control without AQDS in which *M. barkeri* cells encountered standard hydrogenotrophic conditions with CO_2_ as the sole electron acceptor (Figure 1). The 9,10-anthraquinone-2,7-disulfonate disodium salt 98 % (2,7-AQDS) was purchased from TCI 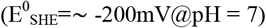 (Huskinson et al. 2014; Wedege et al. 2016) and supplemented to a final concentration of 16 mM (Holmes et al. 2019). All cultures were flushed, and batch fed with a gas mixture of H_2_/CO_2_ (80:20, v/v) at 1.7 bar, stirred at 200 rpm, and incubated at 37 °C.

**Fig. 1.**
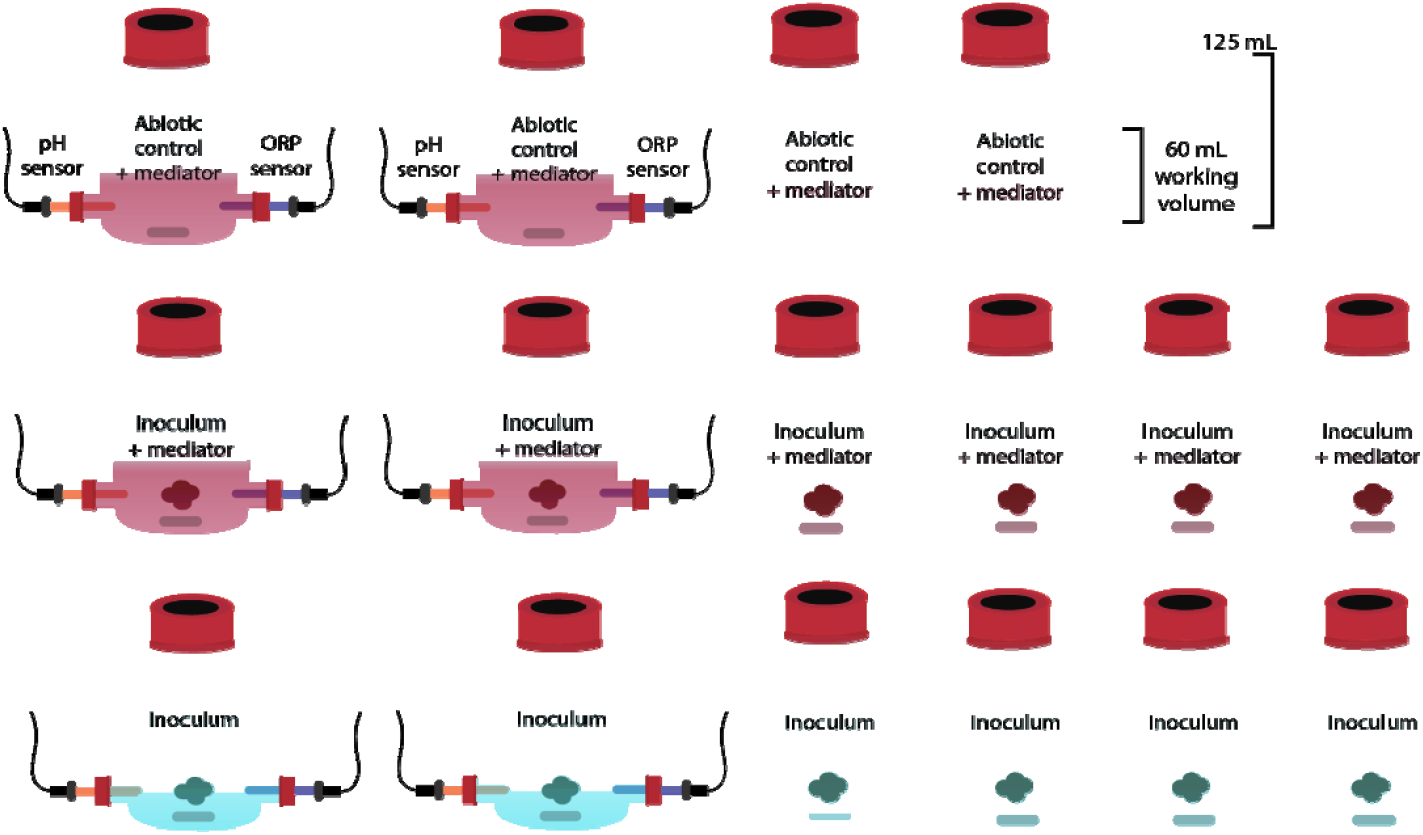
Experimental set-up to evaluate AQDS reduction by *Methanosarcina barkeri*

The experimental setup comprised six replicates per condition, each including two reactors attached to an oxidation-reduction potential (ORP) probe and a pH probe (Atlas Scientific LLC, LongIsland City, USA) for continuous monitoring of redox potential and the redox state (oxidized/reduced) of AQDS. All reactors were sealed with butyl-rubber stoppers and screw caps to uphold pressure and anoxic conditions. The working volume of the reactors was 60 mL, and inoculation was done with 20% of *M. barkeri* pre-culture. Considering that *M. barkeri* cultures required vitamins for growth, the modified 120 DSMZ medium used for the AQDS experiments included the vitamins solution, despite the fact that riboflavin could potentially function as electron shuttle. To ensure anoxic conditions, lower sulfide concentrations (0.3 mM) were used (Holmes et al. 2019), while resazurin and cysteine were omitted.

### 2.3. DNA extractions and qPCR

Liquid samples were collected at the beginning and end of the AQDS reduction experiments for qPCR analysis to assess the cell growth of *M. barkeri* cultures in the presence and absence of AQDS. DNA extractions were carried out using the DNeasy Powerlyzer Powersoil Kit (Qiagen), according to the manufacturer’s instructions. For qPCR analysis, a standard curve for quantification was prepared using an amplification product from a previousl described primer-set (Muyzer et al. 1993). Amplifications were conducted in 25µL reactions using 2xPCRBIO Ultra Mix, 400 nM of each primer and with the following PCR program: 95°C for 1 min, 30 cycles of 95°C for 15 s, 60°C for 15 s, and 72°C for 50 s. The PCR product was cleaned using 0.8x CleanNGS (CleanNA) SPRI beads and the size and quality was assessed using High Sensitivity D1000 on a TapeStation 4150. The product was diluted to 10^8^-10^1^ copies µL^-1^ and used as standard curve. Quantitative PCR (qPCR) assays were performed on samples using Brilliant III Ultra-Fast SYBR® (Agilent) in triplicates. Amplifications were carried out in 20 µL reaction solutions, containing 10 µL 2x Brilliant III qPCR mastermix, 400 nM of each primer, 30nM ROX, 2.7 µL nuclease-free H_2_O, and 5 µL template DNA. The qPCR assay was conducted using the following thermal settings: 95°C for 3 min, followed by 40 cycles at 95°C for 10 s, and 60°C for 15 s, and 95°C for 60 s, 55°C for 30 s, and 95°C for 30 s.

### 2.4. Differential proteomics

Liquid samples were collected from four replicates per condition (from the reactors without the ORP/pH probe) at the beginning and end of the experiment (during the exponential phase). Protein extraction was done using bead-beating and freeze-thawing cycles (Tanca et al. 2015), followed by digestion and purification using the PreOmics iST kit. Proteomic analysis was carried out using a QE Orbitrap system with a 60-minute gradient was used (García-Moreno et al. 2020), and protein annotation was performed against the *Methanosarcina barkeri* 227 UniprotKB reference proteome, with a reversed decoy database using MSFragger (Kong et al. 2017), with variable methionine oxidation. A 0.05 False Discovery Rate (FDR) was employed for quality filtering. To perform differential proteomics data analysis, only proteins retained and detected in 3 out of 4 replicates were used. Initially, quantile-normalization was applied, and any missing values within each group were imputed using the mean of the respective sample group. The log-fold change was calculated by dividing the values of the non-AQDS groups (AQDS-) with AQDS group (AQDS+). Subsequently, *P*-values were computed using a T-test for independent groups, with proteins exhibiting a minimum *p*-value of 0.05 selected for further analysis.

### 2.5. Chemical analyses

Pressure drop in each reactor was monitored using a GoDirect pressure sensor (Vernier) connected to a sterile filter (0.2 µM). The gas composition of the cultures was analyzed using a gas chromatography system (GC-2014, Shimadzu, Japan) equipped with a thermal conductivity detector (TCD) and two different columns. H_2_ utilized a ShinCarbon ST Packed Column (Restek Cat#80486-800, Bellafonte, Pennsylvania, USA) with argon as the carrier gas, while CO_2_ and CH_4_ employed a Porapak Q column (CS-Chromatographie Service GmbH, Manufacturer Item No.: 662, Langerwehe, Germany) with helium as carrier gas.

## 3. RESULTS AND DISCUSSION

### 3.1. Flux of electrons in *M. barkeri*

The capacity of *M. barkeri* to reduce the artificial humic acid homolog 2,7-AQDS was assessed by comparing hydrogenotrophic cultures with and without 2,7-AQDS to abiotic controls (Figure 1). Methane production, H_2_ and CO_2_ consumption, cell growth and proteomic profiles expressed during the reduction of this high-energy electron acceptor were quantified and compared.

Real-time monitoring of 2,7-AQDS reduction was facilitated using an oxidation-reduction potential (ORP) probe, which directly measured the redox state of the culture media in millivolts (mV). Cultures amended with 2,7-AQDS initially exhibited a redox potential of around □ 100 mV vs SHE (standard hydrogen electrode) (Figure 2A). Subsequently, 2,7-AQDS reduction was observed only in reactors inoculated with *M. barkeri*, whereas abiotic controls with identical headspace composition (H_2_/CO_2_, 80:20) showed no redox potential decrease (Figure 2A). This confirms that, despite H_2_ having a lower redox potential than AQDS, biological or chemical catalysts are required for its reduction (Preger et al. 2020). Moreover, the absence of reduction in abiotic controls confirms the role of *M. barkeri* as a catalyst for 2,7-AQDS reduction, consistent with prior observations when 2,6-AQDS was employed as electron carrier for the reduction of Fe(III) oxides (Bond and Lovley 2002; Eliani-Russak et al. 2023).

**Fig. 2.**
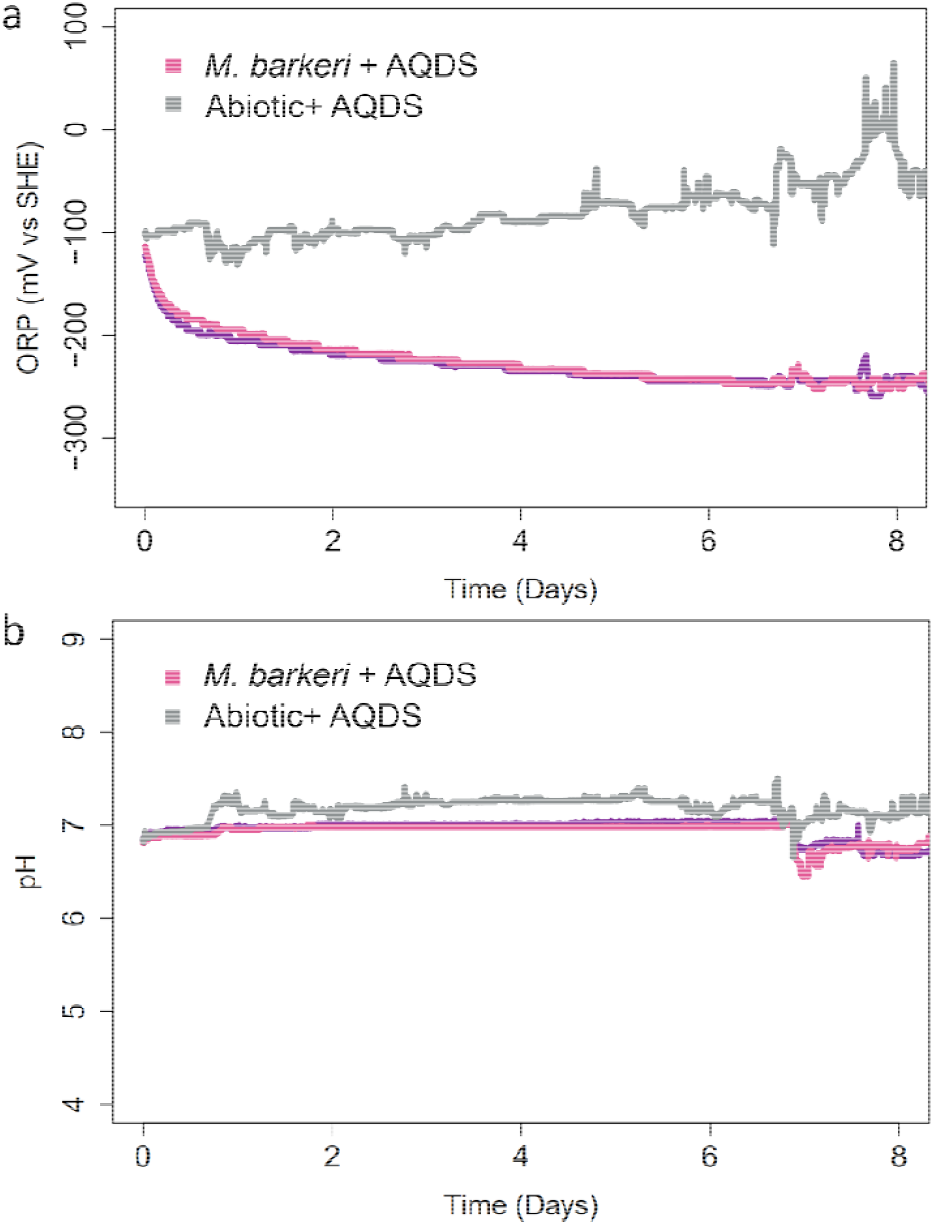
*Methanosarcina barkeri* catalyzed the reduction of 2,7-AQDS under hydrogenotrophic conditions. **a** Oxidation reduction potential measurements (mV vs SHE), **b** pH measurements

Between day 6 to 8, the redox potential of the culture media plateaued around □ 250 mV vs SHE (Figure 2A). Notably, this potential can be influenced by the pH and electrolyte composition of the culture medium. However, throughout the incubation period the microbial reduction of 2,7-AQDS did not influence the pH of the culture medium (Figure 2B). Moreover, the ORP values closely matching the standard redox potential of 2,7-AQDS 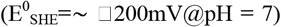 (Huskinson et al. 2014; Wedege et al. 2016) indicated that the reduction process of 2,7-AQDS was incomplete by day 8. By definition, the standard redox potential reflects the state of a half-reaction (Alberty 1998), indicating a 50/50 reduced/oxidized state for the molecule. This ongoing reduction of AQDS underscores the importance of harvesting cultures before reaching the stationary phase for subsequent differential proteomic analyses, a critical consideration for such investigations.

The reduction process of 2,7-AQDS was accompanied by the inhibition of methanogenesis (Figure 3C) and cell growth (Figure 3D), consistent with previous observations during the reduction of Fe(III) oxides via 2,6-AQDS (Bond and Lovley 2002). While the controls without 2,7-AQDS favored methanogenesis through H_2_/CO_2_ conversion (Figure 3A and 3B) for methane and biomass synthesis, cultures amended with 2,7-AQDS appeared t entirely redirect their electron flux from H_2_ towards the higher energy-yielding electron acceptor (Figure 2A). Moreover, the difference in cell biomass likely influenced the extent of H_2_ uptake during 2,7-AQDS reduction (Figure 3A). These findings suggest that although *M. barkeri* possesses the enzymatic machinery to reduce extracellular electron acceptors like 2,7-AQDS with higher redox potential than CO_2_, it lacks the necessary enzymes to conserve energy during this process. The precise mechanism underlying the reduction process remains elusive. I contrast, the related species *Methanosarcina acetivorans* has been shown capable of energy conservation during the reduction of 2,6-AQDS (Holmes et al. 2019) as elaborated in the subsequent section.

**Fig. 3.**
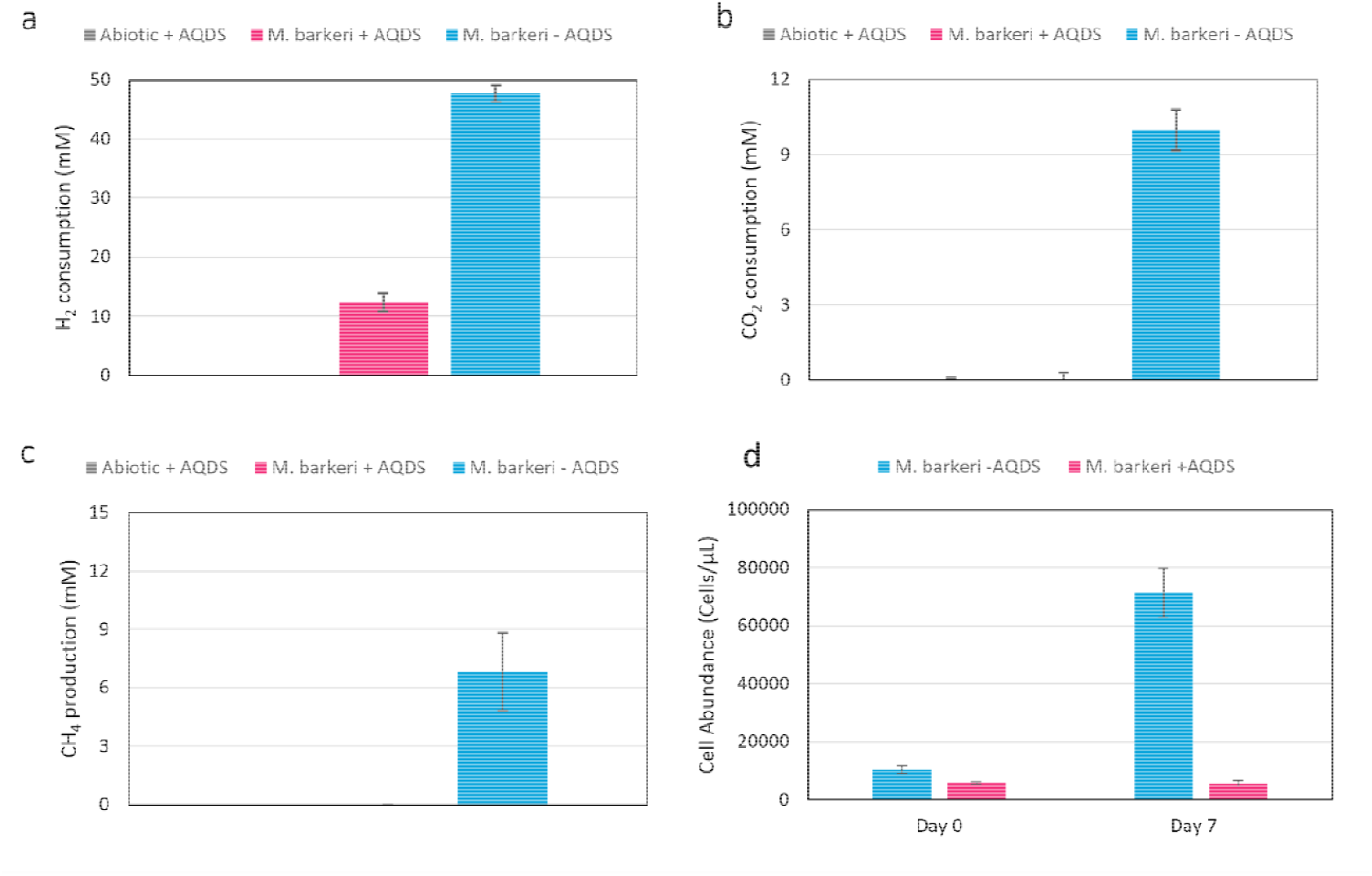
*Methanosarcina barkeri* cultures in the presence and absence of AQDS. **a** H_2_ consumption, **b** CO_2_ consumption, **c** CH_4_ production and **d** qPCR results show the difference between both conditions after 8 days of incubation (before depletion of H_2_ and CO_2_)

### 3.2. Proteomic response from the methanogenesis pathway during 2,7-AQDS reduction

*M. barkeri* and *M. acetivorans* have been extensively studied and are commonly employed as model organisms. Despite belonging to the same genus, they exhibit distinct physiological, ecological, and phylogenetic characteristics, categorizing them into type I and type II *Methanosarcina*, respectively (Zhou et al. 2021). *M. acetivorans* (type II) lacks the energy-converting hydrogenase complex (Ech) and the ability to utilize H_2_ as an electron donor (Sowers et al. 1984; Guss et al. 2009). Instead, it employs the Rnf (Rhodobacter nitrogen fixation) complex as a membrane-bound electron transport chain for ATP production during acetoclastic methanogenesis, along with a seven-heme, membrane-bound, *c*-type cytochrome MmcA to conserve energy for growth via electron transfer to extracellular electron acceptors like 2,6-AQDS (Holmes et al. 2019). Conversely, in type I *Methanosarcina* species such as *M. barkeri*, the observed lack of growth in studies utilizing AQDS (Bond and Lovley 2002) appears related to the absence of the *c*-type cytochrome MmcA (Kletzin et al. 2015). Moreover, *M. barkeri* possesses the Ech hydrogenase complex and the ability to perform hydrogenotrophic methanogenesis (Thauer et al. 2008), relying on intracellular H_2_ cycling for energy conservation and proton motive force generation to support ATP synthesis (Kulkarni et al. 2018; Lovley 2018).

Although energy conservation and cellular growth during quinone reduction have only been demonstrated via the *c*-type cytochrome MmcA in *M. acetivorans*, this reductive process occurs in both natural and artificial environments involving other methanogens. Nevertheless, the cellular mechanism and response to this process remain poorly understood. Consequently, the methanogenic enzymes responsible for the reduction of AQDS and other higher energy-yielding extracellular electron acceptors remain unidentified. Therefore, differential proteomic analyses were performed on *M. barkeri* cultures with and without the addition of 2,7-AQDS, utilizing H_2_ as electron donor.

Figure 4A illustrates a clear response in *M. barkeri* cultures during the reduction of 2,7-AQDS, with distinct sets of proteins being significantly down- and up-regulated. Selection criteria for proteins included a minimum two-fold increase/decrease and a maximum *p*-value of 0.05. Regarding proteins involved in the methanogenic pathway, our findings revealed an upregulation of one of the three hydrogenases required for growth on H_2_/CO_2_ and intracellular H_2_ cycling (Ech, Vht, and Frh) (Mand et al. 2018). Notably, the upregulated membrane-bound Ech hydrogenase complex, comprising 6 different subunits (EchABCDEF), plays a pivotal role in driving the endergonic reduction of ferredoxin (Fd) with a standard potential of approximately □ 500 mV (Bertram and Thauer 1994; Künkel et al. 1998). This reaction is essential for the initial step of methanogenesis from H_2_/CO_2_, namely the reduction of CO_2_ to formyl-methanofuran (CHO-MFR), to occur (Meuer et al. 2002) (Figure 4B). As the Ech hydrogenase is reversible, it can also facilitate the oxidation of reduced ferredoxin (Fd_red_), resulting in both H_2_ generation and the translocation of H^+^ outside the cell (Bott et al. 1986). Given its membrane localization, with two hydrophobic subunits (EchA and EchB) being membrane-bound (Künkel et al. 1998), it is conceivable that among the two upregulated subunits of Ech during 2,7-AQDS reduction, EchB could interact with AQDS (Figure 4C).

**Fig. 4.**
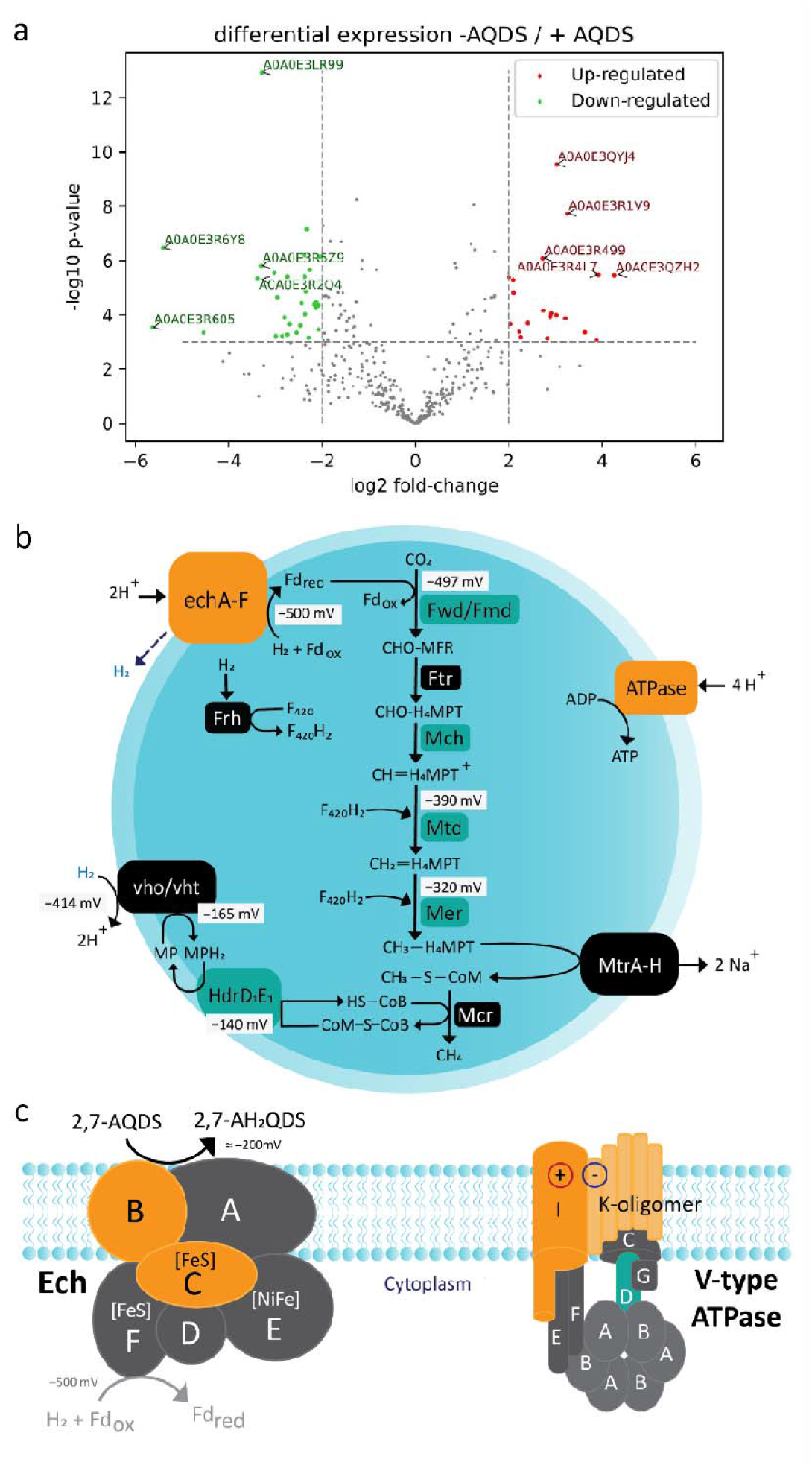
Proteomic response of *Methanosarcina barkeri* cultures (H_2_/CO_2_) during 2,7-AQDS reduction. **a** Volcano plot of the differential proteins expression between 2,7-AQDS reduction versus hydrogenotrophic methanogenesis (without 2,7-AQDS) with a cut-off value of 1.0, **b** *M. barkeri* model shows the upregulated (orange) an downregulated proteins (green) involved in hydrogenotrophic methanogenesis during 2,7-AQDS reduction. Cellular components, methanogenic pathway and standard redox potentials were based on previous models (Thauer et al. 2008; Kulkarni et al. 2018; Mand et al. 2018). **c** Proposed model showing the overexpressed subunits of the membrane bound Ech and V-type ATPase. Electrons from H_2_ are diverted towards AQDS reduction (equation in black) instead of the ferredoxin (equation in grey) due to the higher redox potential of 2,7-AQDS. The enzymes composition and cellular localization were based in previous models (Mulkidjanian et al. 2008; Mand and Metcalf 2019). Abbreviations: Ech, energy conserving hydrogenase; fwd/fmd, formylmethanofuran dehydrogenase; ftr, formyltransferase; mch, methenyl-H_4_MPT cyclohydrolase; mtd, F_420_-dependent methylene-H_4_MPT dehydrogenase; mer, F_420_-dependent methylene-H_4_MPT reductase; mtr, integral membrane methyltransferase; mcr, methyl-coenzyme M reductase; Hdr, the CoB--CoM heterodisulfide reductase; vho/vht, F_420_ non-reducing hydrogenase; Fdox and Fdred, oxidized and reduced ferredoxin, respectively; F_420_ and F_420_H_2_, oxidized and reduced cofactor F_420_, respectively; MP and MPH_2_, oxidized and reduced methanophenazine, respectively

Specifically, in the presence of higher energy-yielding electron acceptors such as 2,7-AQDS, the Ech complex can either catalyze the reduction of ferredoxin at a standard potential of □ 500 mV, enabling subsequent CO_2_ reduction (Bertram and Thauer 1994; Künkel et al. 1998), or reduce 2,7-AQDS at the higher standard potential of –200 mV (Huskinson et al. 2014; Wedege et al. 2016), with the latter reaction being more energetically favorable. As the electron acceptor with the highest redox potential have the strongest affinity for electrons, the electrons divert towards the reduction of 2,7-AQDS instead of generating reduced ferredoxin, (Figure 4C) consequently inhibiting the initial methanogenic step involving CO_2_ reduction. This supports our observation of the absence of CO_2_ conversion during 2,7-AQDS reduction (Figure 3B). Additionally, formylmethanofuran dehydrogenase (fmd), responsible for catalyzing the initial step, was significantly downregulated during 2,7-AQDS reduction (Figure 4B), along with other enzymes involved in subsequent steps of the methanogenic pathway, including methenyl-H_4_MPT cyclohydrolase (mch), F_420_-dependent methylene-H_4_MPT dehydrogenase (mtd), F_420_-dependent methylene-H_4_MPT reductase (mer), and CoB--CoM heterodisulfide reductase (Hdr) (Figure 4B).

Although a prior *in vitro* study involving membranes vesicles from *M. acetivorans* suggested the potential involvement of cytochrome *b* of HdrE in 2,6-AQDS reduction as an electron carrier during Fe(III) reduction (Yan et al. 2018), a separate *in vivo* study in *M. acetivorans* reported slightly lower relative expressions of hdrD and hdrE in AQDS-reducing cells compared to methanogenic cells (Holmes et al. 2019). Moreover, the HdrDE from the *M. acetivorans* MmcA-deficient strain failed to function as the sole AQDS reductase supporting growth during 2,6-AQDS reduction, likely due to its cellular localization in the membrane, as its constituents do not have access to the extracellular milieu *in vivo* (Holmes et al. 2019). Considering this *in vivo* evidence alongside our results showing a significant downregulation of Hdr during 2,7-AQDS reduction, it is improbable that Hdr is the enzyme responsible for AQDS reduction in *M. barkeri*.

Overall, our findings suggest that electrons derived from H_2_ are directed towards the reduction of 2,7-AQDS (via Ech) rather than the methanogenic pathway. Consequently, there is no generation of proton motive force (pmf) to conserve energy for growth through CH_4_ production, as evidenced in our cultures (Figure 3 C, D). Future investigations could clarify whether other energy-converting [NiFe] hydrogenases, like Ech, present in different phylogenetic groups of methanogens, are also implicated in reducing extracellular electron acceptors with higher redox potentials than CO_2_. For instance, there are the two multisubunit membrane-bound [NiFe] hydrogenases, designated Eha and Ehb, found in *Methanothermobacter thermoautotrophicus* and *Methanothermobacter marburgensis* (Tersteegen and Hedderich 1999). From an environmental and biotechnological perspective, understanding how methanogens impact their redox environment by reducing electron acceptors with higher redox potentials in the extracellular milieu is crucial, as this process directly influences microbial metabolisms within anoxic environments.

Furthermore, the significant upregulation of the two membrane-localized subunits, subunit K and I, of the V-type ATP synthase (Mulkidjanian et al. 2008), during 2,7-AQDS reduction, in contrast to the significant downregulation of subunit D (Figure 4C), warrants further investigation into their precise function during 2,7-AQDS reduction, despite the possibility of their up/downregulation being a cellular response to the absence of the proton motive force.

### 3.3. Cellular processes response during 2,7-AQDS reduction

In addition to the proteins associated with the methanogenic pathway, various proteins involved in different metabolic processes were upregulated during the reduction of 2,7-AQDS in *M. barkeri* cultures (Table 1). Regarding redox sensing and cellular response to higher redox potentials, the upregulated proteins included the hemerythrin HHE cation binding domain protein. Although not previously studied in methanogens or Archaea, recent discoveries have shown that bacteria utilize hemerythrin as an oxygen- and redox-sensing domain to adapt to changes in cellular oxygen concentration or redox status, thus maintaining essential physiological processes (Kitanishi 2022). Additionally, rubrerythrin (Rbr), a non-heme iron protein, was upregulated. It serves as an alternative oxidative stress protection mechanism in anaerobic bacteria and archaea by reducing superoxide or hydrogen peroxide (Coulter and Kurtz 2001). The glutaredoxin (GRX) family protein and thioredoxin (TRX) were also upregulated. These ubiquitous redox enzymes play fundamental roles in metabolically and phylogenetically diverse organisms, including serving as reductants for enzymes like ribonucleotide reductase and maintaining cysteine residues of proteins in the reduced state during oxidative stress. They are also involved in redox sensing and regulation of gene expression (Meyer et al. 2009; Ströher and Millar 2012). These findings suggest that during 2,7-AQDS reduction, *M. barkeri* cells sense and regulate their gene expression based on redox potential changes caused by the presence of oxidized 2,7-AQDS, thereby generating the observed proteomic response in the methanogenic pathway.

**Table 1.** Upregulated proteins in *Methanosarcina barkeri* 227 during the reduction of 2,7 AQDS and their associated function.

| Description - Locus ID | ID | fold_<br>change | log2_fold<br>_change | p-value | Associated function |
| --- | --- | --- | --- | --- | --- |
| Mobile element protein MSBR2_0824 | A0A0E3QZH2 | 19.109 | 4.256 | 0.004 | Hypothetical protein |
| Molybdopterin converting factor small subunit MSBR2_3023 | A0A0E3R4L7 | 15.218 | 3.928 | 0.004 | Cofactor biosynthesis |
| Energy-conserving hydrogenase (Ferredoxin), subunit B MSBR2_2764 | A0A0E3R6S7 | 14.734 | 3.881 | 0.046 | Methanogenesis (redox function) |
| V-type ATP synthase subunit K MSBR2_1785 | A0A0E3LQH9 | 12.341 | 3.625 | 0.035 | Energy metabolism |
| V-type ATP synthase subunit K MSBR2_1795 | A0A0E3LQI1 | 12.341 | 3.625 | 0.035 | Energy metabolism |
| HTH iclR-type domain-containing protein MSBR2_0977 | A0A0E3R1V9 | 9.517 | 3.251 | 0.000 | Hypothetical - transcription |
| Uncharacterized protein MSBR2_0643 | A0A0E3QYX0 | 9.233 | 3.207 | 0.021 | Hypothetical protein |
| Cell surface protein MSBR2_0474 | A0A0E3QYJ4 | 8.119 | 3.021 | 0.000 | Unknown function |
| Phosphoribosylformylglycinamidine synthase subunit PurS | A0A0E3LQ36 | 8.049 | 3.009 | 0.018 | Purine biosynthesis |
| Pyridoxamine 5'-phosphate oxidase MSBR2_0873 | A0A0E3QZW4 | 7.519 | 2.910 | 0.017 | vitamin synthesis |
| Uncharacterized protein MSBR2_3222 | A0A0E3R7L4 | 7.394 | 2.886 | 0.019 | Hypothetical protein |
| Carboxymuconolactone decarboxylase family protein MSBR2_2940 | A0A0E3LR60 | 7.091 | 2.826 | 0.044 | Hypothetical protein - aromatic compounds degradation |
| Cytochrome b5 heme-binding domain-containing protein MSBR2_2650 | A0A0E3LR02 | 6.679 | 2.740 | 0.016 | Hypothetical protein - redox function |
| Translation initiation factor 5A | A0A0E3R499 | 6.629 | 2.729 | 0.002 | Translation |
| PRC-barrel domain-containing protein MSBR2_1398 | A0A0E3R175 | 5.264 | 2.396 | 0.025 | Hypothetical protein - translation |
| Translation initiation factor IF-2 subunit beta MSBR2_3427 | A0A0E3R8B0 | 4.783 | 2.258 | 0.042 | Translation |
| Sporulation protein MSBR2_1869 | A0A0E3R4L5 | 4.659 | 2.220 | 0.034 | Hypothetical protein |
| Alcohol dehydrogenase MSBR2_0737 | A0A0E3QZ53 | 4.292 | 2.102 | 0.008 | Redox function |
| ArnR1-like winged helix-turn-helix domain-containing protein MSBR2_2092 | A0A0E3R2T9 | 4.265 | 2.093 | 0.005 | Hypothetical protein - transcription |
| Translation initiation factor IF-2 subunit beta MSBR2_3428 | A0A0E3R872 | 4.097 | 2.034 | 0.026 | Translation |
| V-type ATP synthase subunit I MSBR2_1784 | A0A0E3R469 | 4.011 | 2.004 | 0.005 | Energy metabolism |
| Cell division protein SepF MSBR2_2734 | A0A0E3R6F3 | 3.986 | 1.995 | 0.004 | Hypothetical protein - cell division |
| Uroporphyrinogen-III methyltransferase MSBR2_1699 | A0A0E3R1B8 | 3.308 | 1.726 | 0.046 | Heme synthesis |
| Pyridoxamine 5'-phosphate oxidase putative domain-containing protein MSBR2_1668 | A0A0E3R199 | 3.246 | 1.699 | 0.002 | Hypothetical protein – vitamin synthesis |
| PRC-barrel domain-containing protein MSBR2_2907 | A0A0E3R6N2 | 3.243 | 1.697 | 0.009 | Hypothetical protein - translation |
| Hemerythrin HHE cation binding domain protein MSBR2_1731 | A0A0E3R1D6 | 3.226 | 1.690 | 0.009 | Redox sensing |
| Ruberrythrin MSBR2_2659 | A0A0E3R666 | 3.108 | 1.636 | 0.010 | Oxidative stress |
| Putative snRNP Sm-like protein MSBR2_1127 | A0A0E3R255 | 2.812 | 1.492 | 0.016 | mRNA metabolism |
| Translation initiation factor IF-2 subunit beta MSBR2_3429 | A0A0E3R5H4 | 2.807 | 1.489 | 0.010 | Translation |
| 30S ribosomal protein S7 | A0A0E3R311 | 2.665 | 1.414 | 0.046 | Translation |
| Transcriptional regulator MSBR2_3103 | A0A0E3R7D1 | 2.632 | 1.396 | 0.033 | Transcription |
| Glutaredoxin family protein MSBR2_0595 | A0A0E3LPU1 | 2.416 | 1.273 | 0.002 | Redox function |
| Zinc finger protein MSBR2_1003 | A0A0E3R256 | 2.386 | 1.255 | 0.000 | Cell cycle |
| DUF2795 domain-containing protein MSBR2_3292 | A0A0E3R551 | 2.341 | 1.227 | 0.015 | Hypothetical protein |
| Thioredoxin MSBR2_2931 | A0A0E3R3V9 | 2.279 | 1.188 | 0.001 | Redox function |
| RNA-binding protein YhbY MSBR2_2177 | A0A0E3R4U5 | 2.247 | 1.168 | 0.037 | Translation |
| Energy-conserving hydrogenase (Ferredoxin), subunit C MSBR2_1001 | A0A0E3R2A2 | 2.157 | 1.109 | 0.018 | Methanogenesis (Redox function) |

Additionally, some components related to DNA and RNA processing were upregulated (Table 1). These include translation initiation factor 5A and IF-2, the PRC-barrel domain-containing protein (Anantharaman and Aravind 2002), putative snRNP Sm-like protein, the 30S ribosomal protein S7, zinc finger protein, RNA-binding protein YhbY, and a transcriptional regulator (MSBR2_3103). Furthermore, other significantly upregulated proteins during 2,7-AQDS reduction included the HTH iclR-type domain-containing protein that modulates signal-dependent expression of genes involved in carbon metabolism in bacteria and archaea (Zhang et al. 2002), carboxymuconolactone decarboxylase family protein (MSBR2_2940), required for aerobic growth on aromatic compounds by species in the domain Bacteria (Lessner and Ferry 2007), a hypothetical protein predicted as a cytochrome b5 heme-binding domain-containing protein (MSBR2_2650), uroporphyrinogen-III methyltransferase (HemD-CobA), which is involved in the biosynthesis of siroheme, vitamin B12, heme d1 (Wang et al. 2014), and protoheme (Buchenau et al. 2006), the pyridoxamine 5’-phosphate oxidase (MSBR2_0873, MSBR2_1668), phosphoribosylformylglycinamidine synthase subunit PurS (FGAM synthetase), ArnR1-like winged helix-turn-helix domain-containing protein, cell division protein SepF, a hypothetical sporulation protein (MSBR2_2650), and an alcohol dehydrogenase (Table 1).

On the other hand, aside from the significantly downregulated proteins involved in the methanogenic pathway, our results indicated that ribosomal proteins were also downregulated (See Supplementary Information Table S1), along with the DNA-directed RNA polymerase subunit Rpo1C and Rpo1N, and the replication factor A (SsDNA-binding protein) (Table 2). This suggests that the reduction process of 2,7-AQDS had a negative impact in the enzyme synthesis process by inhibiting proteins related to this process, which can also correlate with the lack of cellular growth.

**Table 2.** Downregulated proteins in *Methanosarcina barkeri* 227 during the reduction of 2,7 AQDS and their associated function.

| Description - Locus ID | ID | fold_<br>change | log2_fold<br>_change | p-value | Associated function |
| --- | --- | --- | --- | --- | --- |
| Methionine--tRNA ligase metG | A0A0E3R6Y8 | 0.024 | -5.405 | 0.002 | Amino acid metabolism |
| Putative (R)-citramalate synthase CimA cimA | A0A0E3R6A0 | 0.043 | -4.540 | 0.035 | Amino acid metabolism |
| F420-dependent methylenetetrahydromethanopterin dehydrogenase mtd | A0A0E3R7X8 | 0.123 | -3.024 | 0.004 | Methanogenesis |
| MarR family protein MSBR2_2434 | A0A0E3R5R9 | 0.126 | -2.988 | 0.040 | Transcription |
| DNA-directed RNA polymerase subunit Rpo1C | A0A0E3R0Y4 | 0.129 | -2.959 | 0.010 | Replication |
| Heat shock protein 60 family chaperone GroEL MSBR2_3399 | A0A0E3R4L2 | 0.138 | -2.857 | 0.040 | Transcription |
| DNA-directed RNA polymerase subunit Rpo1N | A0A0E3R3I3 | 0.143 | -2.807 | 0.020 | Replication |
| V-type ATP synthase subunit D | A0A0E3R475 | 0.149 | -2.744 | 0.038 | Energy metabolism |
| Periplasmic oligopeptide-binding protein OppA MSBR2_1322 | A0A0E3R2X2 | 0.149 | -2.744 | 0.005 | Transport |
| Cell surface protein MSBR2_1960 | A0A0E3LQL4 | 0.154 | -2.701 | 0.026 | Cell surface |
| Methenyltetrahydromethanopterin cyclohydrolase mch | A0A0E3R1A1 | 0.172 | -2.543 | 0.035 | Methanogenesis |
| Dihydroxy-acid dehydratase ilvD | A0A0E3QZZ1 | 0.181 | -2.469 | 0.027 | Amino acid metabolism |
| Acetolactate synthase MSBR2_2700 | A0A0E3LR12 | 0.185 | -2.435 | 0.012 | Amino acid metabolism |
| Chitin binding protein MSBR2_0294 | A0A0E3R0J1 | 0.191 | -2.385 | 0.002 | Cell cycle |
| 3-isopropylmalate dehydrogenase MSBR2_3145 | A0A0E3LRA1 | 0.193 | -2.371 | 0.004 | Amino acid synthesis |
| Rubredoxin-oxygen oxidoreductase MSBR2_2660 | A0A0E3LR04 | 0.195 | -2.358 | 0.018 | Redox fuction |
| Aspartate carbamoyltransferase regulatory chain pyrI | A0A0E3R5C1 | 0.197 | -2.343 | 0.008 | Amino acid metabolism |
| Formylmethanofuran dehydrogenase subunit A MSBR2_1737 | A0A0E3R1E1 | 0.199 | -2.328 | 0.001 | Methanogenesis |
| Phosphate transport regulator MSBR2_0298 | A0A0E3QYJ5 | 0.205 | -2.288 | 0.043 | Hypothetical protein -<br>transport |
| pyruvate, phosphate dikinase MSBR2_1892 | A0A0E3R2B2 | 0.229 | -2.128 | 0.014 | Carbon metabolism |
| 5,10-methylenetetrahydromethanopterin reductase mer | A0A0E3R413 | 0.229 | -2.125 | 0.012 | Methanogenesis |
| 5-methyltetrahydropteroyltriglutamate--homocysteine methyltransferase MSBR2_3431 | A0A0E3R4N3 | 0.237 | -2.079 | 0.013 | Amino acid metabolism |
| 5'-deoxyadenosine deaminase dadD | A0A0E3QY89 | 0.237 | -2.075 | 0.031 | Amino acid metabolism |
| Chitin binding protein MSBR2_1839 | A0A0E3R3P5 | 0.242 | -2.048 | 0.002 | Cell cycle |
| Inosine 5'-monophosphate dehydrogenase MSBR2_0139 | A0A0E3R025 | 0.254 | -1.980 | 0.001 | Purine metabolism |
| Indolepyruvate oxidoreductase subunit IorB MSBR2_1588 | A0A0E3R3Q2 | 0.255 | -1.972 | 0.046 | Redox function |
| Chaperone protein DnaK dnaK | A0A0E3R2B4 | 0.259 | -1.946 | 0.036 | Transcription |
| Chaperonin GroEL | A0A0E3R4E1 | 0.260 | -1.945 | 0.009 | Transcription |
| Serine--glyoxylate aminotransferase MSBR2_2393 | A0A0E3R2U1 | 0.273 | -1.874 | 0.046 | Amino acid metabolism |
| Serine hydroxymethyltransferase glyA | A0A0E3R181 | 0.275 | -1.863 | 0.009 | Amino acid metabolism |
| Phosphoserine phosphatase MSBR2_3301 | A0A0E3R7J6 | 0.282 | -1.825 | 0.001 | Amino acid metabolism |
| DUF169 domain-containing protein MSBR2_1542 | A0A0E3R1M3 | 0.292 | -1.774 | 0.013 | Hypothetical protein |
| S-layer family duplication domain-containing protein MSBR2_2461 | A0A0E3R3R4 | 0.308 | -1.699 | 0.033 | Hypothetical protein |
| Formylmethanofuran dehydrogenase subunit A MSBR2_1990 | A0A0E3LQM0 | 0.317 | -1.657 | 0.008 | Methanogenesis |
| Imidazole glycerol phosphate synthase subunit HisF | A0A0E3R4S1 | 0.320 | -1.642 | 0.020 | Amino acid metabolism |
| DUF169 domain-containing protein MSBR2_3044 | A0A0E3R6X0 | 0.326 | -1.617 | 0.013 | Hypothetical protein |
| Exosome complex component Rrp41 | A0A0E3R0S8 | 0.331 | -1.597 | 0.001 | RNA/ protein degradation |
| Phosphoribosylaminoimidazole-succinocarboxamide synthase purC | A0A0E3R6X6 | 0.340 | -1.555 | 0.008 | Purine metabolism |
| formylmethanofuran dehydrogenase MSBR2_1734 | A0A0E3R423 | 0.351 | -1.510 | 0.016 | Methanogenesis |
| S-inosyl-L-homocysteine hydrolase MSBR2_0325 | A0A0E3LPN7 | 0.359 | -1.480 | 0.033 | Amino acid metabolism |
| Chitin binding protein MSBR2_2060 | A0A0E3LQN4 | 0.360 | -1.475 | 0.032 | Cell cycle |
| valine--tRNA ligase MSBR2_1685 | A0A0E3LQF9 | 0.361 | -1.469 | 0.023 | Translation |
| Heat shock protein 60 family chaperone GroEL MSBR2_0246 | A0A0E3R0B0 | 0.372 | -1.425 | 0.020 | Transcription |
| Cell division protein FtsZ | A0A0E3LRA0 | 0.391 | -1.353 | 0.030 | Cell cycle |
| (5-formylfuran-3-yl)methyl phosphate synthase mfnB | A0A0E3R7Y4 | 0.401 | -1.320 | 0.040 | Methanogenesis |
| Cell surface protein MSBR2_1341 | A0A0E3R2Z1 | 0.406 | -1.300 | 0.028 | Cell surface |
| CoB--CoM heterodisulfide reductase 2 iron-sulfur subunit D MSBR2_2817 | A0A0E3R4B8 | 0.415 | -1.271 | 0.011 | Methanogenesis |
| Aminotransferase MSBR2_2390 | A0A0E3LQV0 | 0.417 | -1.262 | 0.012 | Amino acid metabolism |
| PAC2 family protein MSBR2_3295 | A0A0E3LRD1 | 0.418 | -1.259 | 0.000 | RNA/ protein degradation |
| formylmethanofuran dehydrogenase MSBR2_1989 | A0A0E3R476 | 0.450 | -1.152 | 0.003 | Methanogenesis |
| Replication factor A (SsDNA-binding protein) MSBR2_0036 | A0A0E3QXY2 | 0.453 | -1.141 | 0.002 | Replication |

## 4. Conclusion

In anoxic environments, both natural and man-made, methanogenesis thrives under extremely low redox conditions to facilitate the reduction of CO_2_. CO_2_ stands out as one of the least favorable electron acceptors utilized in anaerobic metabolisms. However, in anoxic environments methanogens encounter other electron acceptors with higher redox potentials, such as iron(III) oxides and quinones. Using *Methanosarcina barkeri* as a model methanogen, this study aimed to understand the physiological and proteomic response during the reduction of the artificial quinone 2,7-AQDS. The presence of higher energy-yielding electron acceptors directly impacts methane production not only in natural environments but also during methanogenesis in biotechnological processes.

Our findings confirm that during the reduction of 2,7-AQDS, *M. barkeri* cultures did not conserve energy for cellular growth. Instead, the flux of electrons diverted towards 2,7-AQDS rather than flowing towards the reduction of ferredoxin, a crucial reaction involved in the initial methanogenic step to reduce CO_2_ into formyl-methanofuran. As a result, the electrons derived from H_2_ flowed towards the reaction with higher redox potential (2,7-AQDS reduction), leading to the non-utilization of CO_2_ and consequent inhibition of methanogenesis, resulting in a lack of methane production and cellular growth.

The impact on the methanogenesis pathway was further supported by the set of significantly upregulated and downregulated proteins. During 2,7-AQDS reduction, the membrane-bound subunits B and C of the energy-conserving hydrogenase Ech were upregulated, while enzymes involved in the subsequent steps of methanogenesis were downregulated. This underscores that higher energy-yielding electron acceptors such as 2,7-AQDS directly inhibit hydrogenotrophic methanogenesis in the type I *Methanosarcina*, like *M. barkeri*, which lack membrane-bound multiheme *c*-type cytochrome (MmcA).

Further studies on the function of Ech subunits B and C will be essential to elucidate how electrons are precisely transferred to 2,7-AQDS, and whether the two Ech homologs, Eha and Ehb, present in other phylogenetic groups of hydrogenotrophic methanogens, are responsible for quinone reduction processes. Ultimately, the reduction processes of higher energy-yielding electron acceptors and the increase of redox potential during methanogenesis have significant implications from both environmental and biotechnological perspectives.

