## Supplementary material for "Elucidating redox-driven inhibition of methanogenesis by an artificial quinone in *Methanosarcina barkeri*: Integrated proteomic and physiological evidence": suplementary 1

**Supplementary information**

**S1.** Downregulated ribosomal proteins in *Methanosarcina barkeri* 227 during the reduction of 2,7 AQDS

| Description - Locus ID | ID | fold_change | log2_fold_change | *p*-value |
| --- | --- | --- | --- | --- |
| 50S ribosomal protein rpl29 | A0A0E3R605 | 0.020 | -5.627 | 0.029 |
| 30S ribosomal protein S3Ae | A0A0E3R2Q4 | 0.096 | -3.383 | 0.005 |
| 30S ribosomal protein S5 | A0A0E3R5Z9 | 0.101 | -3.307 | 0.003 |
| 50S ribosomal protein L10 | A0A0E3LR99 | 0.102 | -3.292 | 0.000 |
| 30S ribosomal protein S19e | A0A0E3R4I7 | 0.208 | -2.268 | 0.003 |
| 30S ribosomal protein S3 | A0A0E3LQY6 | 0.221 | -2.179 | 0.012 |
| 50S ribosomal protein L6 rpl6 | A0A0E3R3X1 | 0.265 | -1.914 | 0.009 |
| 30S ribosomal protein S4 | A0A0E3LQY1 | 0.329 | -1.603 | 0.018 |
| 50S ribosomal protein L23 | A0A0E3R372 | 0.399 | -1.325 | 0.020 |
